# Reproductive phenology of the eastern oyster, *Crassostrea virginica* (Gmelin, 1791), along a temperate estuarine salinity gradient

**DOI:** 10.1101/2022.11.21.517094

**Authors:** Kaili M. Gregory, Katherine McFarland, Matthew P. Hare

## Abstract

Low salinity can negatively affect reproduction in estuarine bivalves. The spatial and temporal extent of these effects are important to inform models of population dynamics, environmental risk assessments, restoration efforts, and predictions of climate change effects. We hypothesized that oysters at low salinity sites would have delayed gametogenesis compared to their higher salinity counterparts in downstream experimental cages. The timing of gametogenesis and spawning was observed June – August for 2-year-old oysters from three distinct ancestries (Native, Hatchery, Aquaculture), outplanted at age 1 month along the salinity gradient (3–30 psu) of a temperate estuary. A second season of data was collected from 3-year-old Aquaculture oysters (comparable to year 1 data) and Native adult oysters transplanted one year prior. Dermo was very low both years. A delay in gametogenesis and spawning was observed for all ancestries at low salinity relative to higher salinity sites during July and August of the first year but not the second year. In contrast, June showed the reverse pattern with northern low salinity sites having more advanced gonad index (2.65) than a high salinity site (1.46). This difference in average gonad index was 2.65 vs 1.46, respectively, for the Native line and 2.62 vs 2.08 for Aquaculture. Low salinity seemed to not only induce earlier gametogenesis in June, but also extended the reproductive season relative to higher salinity sites. Among oyster ancestries, the Aquaculture line stood out as having 30 – 48% lower gametogenic synchrony within sites, but only in 2018. Despite some dependence of reproductive phenology on salinity variation, the Native low salinity population demonstrates notable reproductive plasticity in the completion of a reproductive cycle across a wide range of salinities, an encouraging result for potential future restoration strategies.

## INTRODUCTION

Estuaries are ecologically vital ecotones characterized by a salinity gradient (Telesh and Khlebovich 2010). Average salinity levels fluctuate with tide, weather, river discharge, and season, with a general rise in salinity as proximity to the ocean increases (Warner et al. 2005). Many species are adapted to the variable salinity conditions of estuaries, whereas others are interlopers that take advantage of estuarine habitats during one or more life stages (Able 2005; Lellis-Dibble 2008). Among estuarine adapted species, physical niche is often described using a habitat suitability index constructed from viability or growth rate across different combinations of estuarine environments (Barnes et al. 2007; Swannack et al. 2014; Linhoss et al. 2016). These indices help guide habitat protection priorities and restoration planning by modelling how species would fare in specified areas. There are two aspects of population variation that are rarely accounted for with these indices: (1) phenotypic plasticity and (2) genetic differences among local or supplemented populations. Phenotypic plasticity can dramatically broaden the realized niche of a population, so experiments that inform performance indices should ideally include a full range of acclimation effects to accurately predict performance at the margin.

Genetic differences among local populations, originating from within-generation selection rather than multigenerational adaptation (Marshall et al. 2010; Sanford and Kelly 2011), have potential to affect population resilience through variation in life history traits, including the degree of phenotypic plasticity (Eierman and Hare 2015).

With metapopulation structure as our within-estuary model, and given the added productivity and resilience expected from portfolio effects when component populations have somewhat independent life history or population dynamics (Lipcius et al. 2008; Schindler et al. 2010), it is important to document performance variation among the diverse habitats within an estuary. Here, we focus on a trait that is poorly known for many estuarine organisms - gametogenic and spawning phenology relative to the salinity gradient. Spawning phenology differences across an estuarine metapopulation could have a large impact on recruitment and population dynamics because the timing of larval production, relative to hydrodynamic, microalgal, and community ecology dynamics, can be expected to constrain or promote larval survivorship (Starr et al. 1990; Morgan 1995).

Our study organism is the eastern oyster (*Crassostrea virginica*, Gmelin, 1791) because of its keystone role, contributing valuable ecosystem services to estuarine communities such as water filtration, benthic pelagic coupling, and a structurally complex benthic reef habitat used by many commercial fish species (Coen et al. 2007; Beck et al. 2011; Bricker et al. 2020; Rose et al. 2021). The synergistic mix of coastal degradation, overharvesting, disease, eutrophication, and climate change threaten to degrade oyster populations to functional extinction in some regions, if it has not done so already (Beck et al. 2011). While salinity is considered to be a weak or negligible environmental factor for triggering spawning in eastern oysters relative to temperature and chemical cues (Thompson et al. 1996), its effect on pre-spawning gametogenic development and overall reproductive timing is much less studied. More fully understanding the effect of salinity variation on reproductive phenology has the potential to inform eastern oyster population dynamic models, environmental risk assessments, restoration efforts, and improve predictions of the effects of climate change (Barnes et al. 2007; Marshall et al. 2010; Levinton et al. 2011). For these reasons, this study primarily examines the effect of natural estuarine salinity variation on the reproductive phenology of the eastern oyster.

Unfortunately, many estuaries no longer have extant populations of eastern oysters along the entire salinity gradient. In fact, the desire to restore full metapopulation structure within estuaries provides an important motivation for understanding how salinity variation affects phenology. The effects of salinity variation on overall reproductive timing has received little attention outside of aquaculture settings. It is well-established that the timing of spawning for eastern oysters is closely correlated with rising temperatures, particularly in temperate estuaries where reproduction has a more pronounced seasonality (Loosanof and Davis 1952; Davis and Calabrese 1964; Bourlès et al. 2009). The interactive effects of salinity and temperature influence multiple aspects of oyster life history traits such as mortality, growth, and survival at all life stages (Davis and Calabrese 1964; Rybovich et al. 2016; McFarland et al. 2022). Multiple studies have shown that low salinity environments (< 6 psu) can inhibit or depress gametogenesis in oysters (Butler 1949; Loosanof 1953; Shafee and Daoudi 1991; Shumway 1996; Honig et al. 2014; Volety et al. 2017), but the effect of salinity variation specifically on reproductive phenology has been much less studied. Butler (1949) addressed how natural salinity variation affects gametogenic progression using histology, yet that case involved unusually protracted and extreme low salinity conditions in the upper Chesapeake Bay rather than the average salinity gradient experienced by an estuarine oyster metapopulation. The strongest data relating salinity to reproductive phenology that we are aware of come from a 15-year time series of monthly oyster collections at five sites along the Caloosahatchee River Estuary, Florida, ranging from 12 to 28 average salinity in September (the annual salinity low point). Based on gonad index averages over 15 years for wet, moderate, and dry subsets, McFarland et al. (2022) found that the upper-most site had the earliest gametogenic development and the longest overall reproductive season. Dermo disease was negatively correlated with salinity and average gonad index, making it difficult to infer the separate phenology effects of these two factors (McFarland et al. 2022).

Dermo disease is caused by the pathogen *Perkinsus marinus*. Moderate to heavy infection reduces an oyster’s investment in reproduction and shell growth, with the potential to cause mortality and devastate oyster populations (Chu and La Peyre 1993; Powell et al. 1996). This disease poses a pervasive risk to populations across the eastern seaboard and is thus of great concern to management and restoration planning (Andrews 1988; Dittman et al. 2001; Mackenzie 2007). Non-lethal infection intensities can have significant negative effects on gonad size and gametogenic development (Dittman et al. 2001). Dermo disease risk is largely limited to high and moderate salinity environments (≥12 psu) because of the pathogen’s environmental limitations (Powell et al. 1996; Levinton et al. 2011). Thus, while oyster gametogenesis is inhibited in extreme low salinity environments, such areas can also serve as a refuge from disease (La Peyre et al. 2003; La Peyre et al. 2016). Documenting Dermo prevalence can not only help to interpret potential mechanisms limiting gametogenesis between high and low salinity sites, but it also provides a critical baseline metric to inform restoration planning and monitoring the health of restored populations.

The Hudson River Estuary (HRE) provides a particularly interesting case study. This estuary includes multiple named water bodies in the brackish zone above and below New York City (Fig. 1). The HRE was once home to a highly abundant, economically valuable population of eastern oysters (Franz 1982; Kurlansky 2006). Overharvesting, water pollution, habitat destruction, and urbanization led to a precipitous decline of the population in the early 20^th^ century (Kurlansky 2006; Mackenzie 2007). Today, a self-sustaining eastern oyster population is largely absent from the HRE except for a small remnant population in the Tappan Zee-Haverstraw Bay (TZHB) portion of the estuary near Irvington, NY (Fig. 1), where average salinity is low (3-12 psu) (Medley 2010; Starke et al. 2011). Recent unpublished data (Hare) suggest that the TZHB population, while reproducing annually and self-recruiting consistently (Levinton et al. 2011; McFarland and Hare 2018; AKRF Inc. et al. 2021), is barely contributing any spat recruitment downstream that could help recover a self-sustaining population in lower parts of the estuary where water quality has improved and restoration efforts abound (Stinnette et al. 2018; McCann 2019). Based on published habitat suitability indices for *C. virginica*, the persistence and reproductive output of the TZHB population at very low salinities is enigmatic but provides hope that when restoration broodstock are needed, local native oysters will be considered a viable option. Our goal was to test the spawning capacity of TZHB oysters when transplanted to other parts of the estuary, and measure salinity effects on gametogenic timing.

**Fig. 1.**
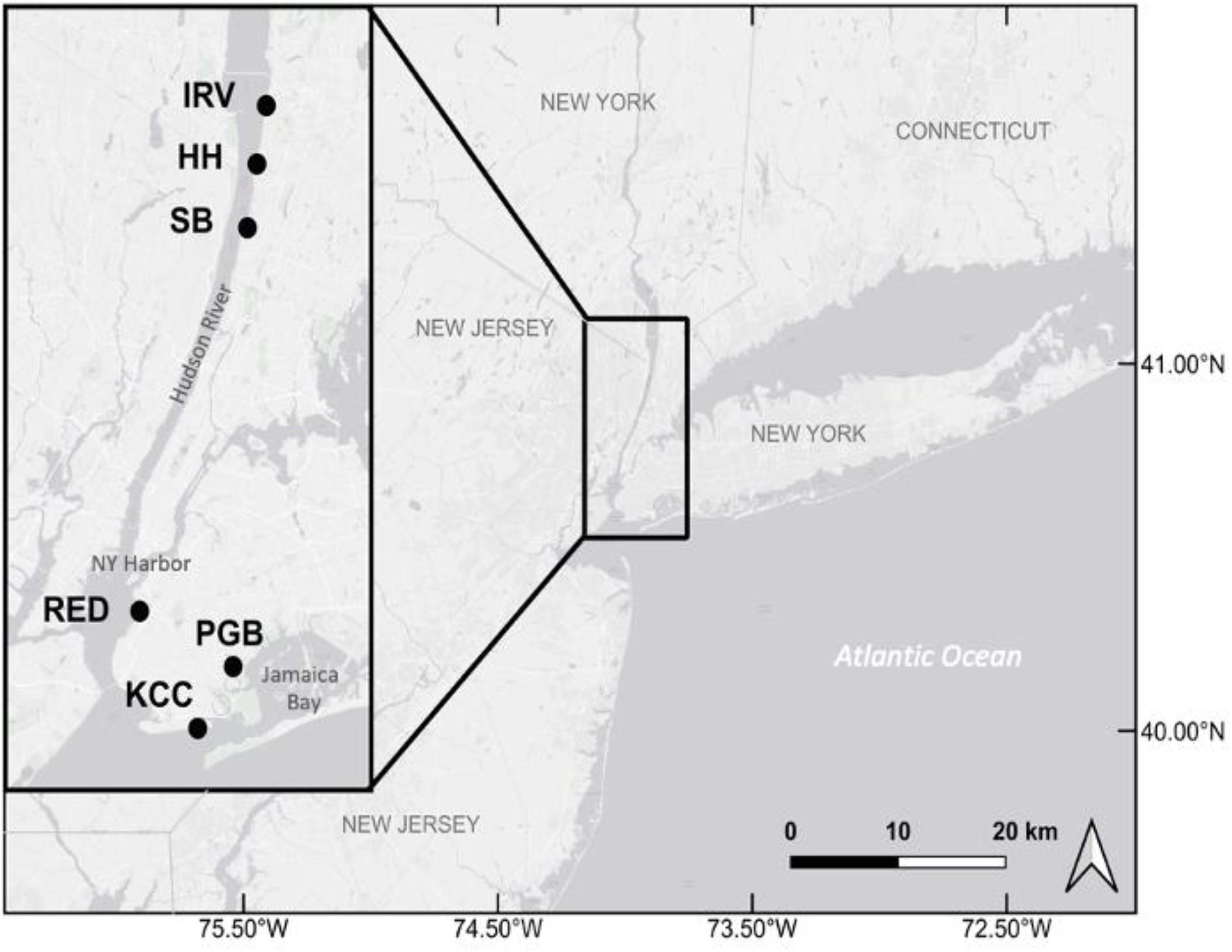
Map of sampling locations on the Hudson River Estuary in New York, United States. Sampling locations are in three named regions of the estuary: Irvington Boat Club (IRV), Hastings on the Hudson (HH), and Yonkers Science Barge (SB) are north of New York City in the “Lower Hudson River”, also known as the North River; Redhook (RED) is in New York Harbor; Paerdegat Basin (PGB), and Kingsborough Community College (KCC) are both in Jamaica Bay.

To increase the generality of this study we compared gametogenic phenology of oysters from three population sources. Choosing an oyster source for restoration typically requires evaluation of potential trade-offs with respect to restoration efficacy, involving logistics, availability, cost, genetic diversity and relative fitness in the specific restoration environment (Hornick and Plough 2019; Hornick and Plough 2022). There currently are no relative performance data available on gametogenesis and phenology for oysters with distinct ancestries, even though these phenotypes can vary due to either local adaptations (genetic factors) or acclimation to distinct environments (epigenetic factors). Local native transplants (source 1: “Native”), recommended to maximize genetic diversity while minimizing maladaptive gene flow (Camara and Vadopalas 2009), involve transplants from naturally recruiting populations within the same estuary or region. For simplicity we will refer to naturally recruited oysters as “Native”, acknowledging that they are not necessarily pristine or genetically non-admixed with aquaculture lines at present or in the past. Native transplants can potentially be done at any post-settlement life stage, and do not involve hatchery propagation. Hatchery propagated stocks for restoration or supplementation can derive from wild broodstock spawned in the hatchery (source 2: “Hatchery”) or from artificially selected lines used in aquaculture (source 3: “Aquaculture”). Most U.S. eastern seaboard hatcheries producing oysters sell primarily to aquaculture growers and do not typically work with wild broodstock, whereas restoration hatcheries and some research hatcheries spawn and propagate wild oysters for restoration (Hornick and Plough 2019). Thus, there can be many practical trade-offs affecting the decision to use one oyster source or another for restoration. A recent survey of New York/New Jersey projects in which oysters were seeded on restored benthic habitat revealed that 19 out of 20 had obtained seed from an out of state hatchery, with at least partial use of local broodstock in only 3 cases (McCann 2019).

Here, we compare the timing of gametogenic development and among-oyster synchrony for these three oyster ancestries along the salinity gradient in a temperate estuary. By quantifying gonad development at locations ranging from 3 to 30 psu, we test for the effects of salinity and ancestry on reproductive phenology. Our first hypothesis is that gametogenesis is delayed in the early-reproductive season (June) at sites with lower salinity relative to the higher salinity sites (Butler 1949; Shafee and Daoudi 1991; Shumway 1996; Honig et al. 2014; Volety et al. 2017).

Comparing oyster ancestries, our hypothesis is that the Native line, from the local TZHB population, will demonstrate the most distinct phenology out of the three lines. More specifically, the long history of the TZHB population at an isolated, low salinity location (McFarland and Hare 2018), leads us to hypothesize that local adaptation or acclimation has reduced gametogenic suppression at low salinity, compared to the Aquaculture and Hatchery lines, both of which involved hatchery-produced seed (juveniles) from moderate salinity broodstock. Accordingly, the gametogenic delay predicted by hypothesis 1 is only expected in these latter two lines.

## METHODS

### Study Species

Eastern oysters (hereafter, oysters) are protandrous hermaphrodites, maturing first as males and later becoming females, and spending some period of the winter, depending on temperatures, with undifferentiated sex (Kennedy and Battle 1964). After sex differentiation, oysters begin gametogenesis, building and developing gonad tissue and gamete cells (eggs or sperm). The oyster is a synchronous spawner, triggered by temperatures greater than 20°C to release gametes into the water column as nearby individuals do the same (Loosanof and Davis 1952; Kennedy and Battle 1964; Barber et al. 1991; Barber 1996; Mann et al. 2014). In temperate waters like the HRE, the reproductive period (gametogenesis and spawning) begins in the Spring and ends in the Fall. As it gets colder, oysters sometimes achieve a smaller fall spawn (Hayes and Menzel 1981) and then enter a quiescent stage in the winter during which gonads are dormant and mostly undifferentiated until the next spring (Kennedy and Battle 1964; Hayes and Menzel 1981; Mann et al. 2014).

### Outplant locations

During a previous study initiated in 2016, juvenile oysters were deployed in cages along the estuarine salinity gradient to measure and compare survivorship and growth over 3 years. For this study we sampled lines from six of these outplant locations in both 2018 and 2019 (Fig. 1). The locations at Irvington Boat Club (IRV), Hastings on the Hudson (HH), and Yonkers Science Barge (SB) represent the low salinity region of the HRE (3-13 psu). The Redhook (RED) location represents an intermediate salinity (15-25 psu) in New York Harbor. Paerdegat Basin (PGB) and Kingsborough Community College (KCC) are both in Jamaica Bay (connected to the lower HRE) and have the highest salinities (20-30 psu) (Fig. 1). Temperature and salinity were recorded as point data during each sampling trip at each location, along with semi-continuous measurements collected using YSI 600 OMS (YSI Inc. Yellow Springs, Ohio) sondes deployed at KCC and RED. Environmental data for HH, IRV and SB were obtained from Riverkeeper Incorporated’s Water Quality Program in the Hudson River Estuary (https://www.riverkeeper.org/water-quality/hudson-river/), the Hudson River Environmental Conditions Operating System (HRECOS) location at Piermont Pier (across the river from IRV), and the Center for the Urban River at Beczak (sonde deployed at SB) (http://hudson.dl.stevens-tech.edu/hrecos/d/index.shtml). Water quality data for PGB were obtained from a Harbor Water Quality dataset provided by the New York City Department of Environmental Protection (https://data.cityofnewyork.us/Environment/Harbor-Water-Quality/5uug-f49n).

### Oyster ancestry

At each location, three distinct oyster lines were compared: Native, Hatchery, and Aquaculture. The 2018 Native line was obtained in August/September 2016 as natural spat recruits on bivalve shell deployed in the vicinity of the TZHB remnant population, then transplanted as 1 month old spat to various cage grow-out locations (Fig. 1). The Hatchery line represents wild genetic diversity from a moderate-salinity population with standard hatchery procedures imposing some genetic bottlenecking (Hornick and Plough 2019). The Hatchery line was produced by strip-spawning 12 males and 12 females collected from a native population in Edgartown Great Pond, a moderate-salinity (15-25 psu) lagoon in Martha’s Vineyard, Massachusetts. The larvae were then cultured by the Martha’s Vineyard Shellfish Group and sent to the Cornell Cooperative Extension hatchery in Southold, New York, to set on micro-clutch and grow out in a nursery upwelling system for three weeks during August 2016 before deploying to sites across the HRE (Fig. 1). Larval culture was at 27-28 psu and the nursery was 28 psu. Finally, a moderate-salinity aquaculture line was acquired as seed oysters (clutch-less spat) in August 2016 for outplant in the same cages. We estimated that all three lines were within approximately 2-3 months of the same age and were thus approximately two years old when sampled in 2018.

The 2019 samples consisted of the same Aquaculture line (now 3 years old) and TZHB native oysters dredged as adults (mean length = 74.57*±* 1.13mm) in June 2018. The dredged samples were immediately transplanted to experimental cages at the study sites (Fig. 1). Unlike the 2018 TZHB native oysters that developed from spat in the experimental cages at each study site (nearly full post-settlement opportunity for developmental plasticity), 2019 TZHB native oysters had acclimated as adults for 1 year at each study site before summer sampling in 2019. The 2019 Native cohort is recognized and considered a fourth treatment group because of their alternate collection and deployment strategy. Due to common ancestry, the 2019 cohort is still included in the discussion of ancestry effects on salinity.

### Oyster sampling and histology preparation

Oysters were collected and sampled for histology three times during the summer of 2018 (June 15-21, July 8-13 and August 8-13). Logistical and financial constraints limited collections to two dates in 2019 (July 28-August 1 and September 1-4), and Aquaculture individuals were not available at every location in 2019. The number of individuals that could be histologically analyzed was limited by oyster survivorship as well as funding. At each sampling site visit, shell height was measured to the nearest millimeter using Mitutoyo Calipers (Mitutoyo America Co., Aurora, IL, USA) from the hinge to the farthest edge of the shell prior to dissections.

In 2018, oysters were transported live in coolers with ambient HRE water to the Fort Totten Urban Field Station Laboratory in Queens, New York, (US Forest Service) where dissection and fixation of tissue sections was performed. In 2019, oysters were transported back to Cornell University live in coolers with ice packs, location specific HRE water, and aeration for processing the next day. After cleaning and shucking the oysters, a standardized 4mm cross section was taken to include reproductive tissue at the intersection of the labial palps and gills (Morales-Alamo and Mann 1989; Howard et al. 2004). The tissue cross section was placed in a labeled plastic cassette and preserved in Davidson’s Fixative for 7 days and then 70% ethanol (Fisher et al. 1996). Samples were then sent to the Cornell University Animal Health Diagnostic Center for histology slide preparation with hematoxylin and eosin staining.

### Categorical gametogenesis analysis

A common approach to measuring oyster reproductive status is with qualitative categorical scoring of gametogenesis progression from histological sections (Fisher et al. 1996). We scored the 5 stages defined by Lango-Reynoso et al. (2000), Barber (1996), and Kennedy & Battle (1964) (Table 1). Histology slides were examined with a compound microscope under 100x total magnification to determine sex and gonad developmental stage. Gametogenesis was scored by KMG. A subset of oysters with borderline category assignments were also scored independently by MPH until consistent agreement was obtained.

**Table 1.** Descriptions of gametogenic development stages for *Crassostrea virginica* (Kennedy and Battle 1964; Barber 1996; Lango-Reynoso et al. 2000)

| Stage Number | Stage Title | Description |
| --- | --- | --- |
| 0 | Inactive | Follicles are nonexistent or elongated, with walls consisting of undifferentiated germinal epithelium. Sex cannot be determined |
| 1 | Early-active | Follicles contain oogonia or spermatogonia and primary oocytes or spermatocytes (no free oocytes or spermatozoa) |
| 2 | Late-active | Secondary (free) oocytes and spermatocytes predominate in the follicles; there are some spermatozoa |
| 3 | Mature | Mature gametes (ova or spermatozoa) totally filling the follicles; presence of ova with distinct nucleus and nucleolus, spermatozoa oriented with tails toward the follicle lumen |
| 4 | Spawned | Follicles have gaps devoid of gametes, although numerous gametes may still remain, follicle walls may be broken. Redevelopment as evidenced by increased number of primary oocytes or spermatocytes |
| 5 | Reabsorbing | Follicles have a shrunken appearance and contain numerous phagocytes and products of reabsorption; gametes are refractory, and development is not evident |

### Dermo disease testing

To test for the presence of *P. marinus* and measure intensity of infection, a small section of rectum tissue and approximately 0.5 cm^2^ from the posterior gill tissue were removed from the same individuals on which histology was performed and placed in a culture tube containing 9.5 mL of Ray’s Thioglycollate Medium (RFTM; (Ray 1952; Ray 1954). Then, 0.5mL antibiotic solution (equivalent to 500 units of penicillin G and 500 units of streptomycin per mL of medium) and 50 μL of nystatin solution were added to the culture tube. The cultures were incubated at 25-26 °C in the dark for 5-7 days. After the incubation period, all tissues were removed from each tube and placed on a clean glass microscope slide. The tissue was macerated using a razor blade and 3-4 drops of Lugol’s iodine was added before applying a cover slip. Each slide was immediately examined at 100-400x magnification for Dermo cells using the Mackin scale of 0 to 5, with 0 indicating no infection and 5 heavily infected (Mackin 1962; Dittman et al. 2001). Because infections were so sparse, several slides were sent for confirmatory readings by Iris Burt at Rutgers Haskin Shellfish Research Lab, New Jersey. All individuals sampled in 2019 for histology (n = 240) were tested for levels of *P. marinus* infection.

### Statistical Analyses

The proportions of each gametogenic stage were calculated for each unique combination of location, month, and line. A set of ordinal logistic models were created to test the relationship between proportions of each gonad development stage and location, month, sex, and line for the 2018 sampling year (Table 2). The moderate-salinity site (RED) had extremely low cumulative survivorship (by August 2018: Native-3%, Hatchery-0%, Aquaculture-9%), resulting in a small sample size, so it was not included in the model. Given that complete or nearly complete gametogenic data were available for sites representing two environmental extremes in the estuary, northern low salinity and southern high salinity, location was used as a proxy for salinity and included in all models. Month was included in all models because observations were made at this time interval during the typical main gametogenic season for oysters at this latitude (Kennedy and Battle 1964). All resulting possible interaction effects were included for 2018 models. The 2019 model did not include a line variable because of limited data across ancestries, and because the Native line was based on adult transplants, not juvenile transplants like in 2018. Instead, 2019 Aquaculture data are described qualitatively, and only Native data were included in a location + month + location*month model to best mirror the top ranked model in the 2018 model selection (Table 2). Models were ranked in terms of AIC values with the criterion that well supported models have a delta AIC value < 2 (Burnham and Anderson 2002). Pairwise *post hoc* contrasts were performed using the ‘emmeans’ package (Lenth 2022). All statistical analyses were performed using R and RStudio (R Core Team 2018).

**Table 2.** Results of 2018 model selection of ordinal logistic models for gametogenic development stages. Models are ranked in terms of AIC values

| Model # | Description | Df | Log Likelihood | AIC | Delta AIC | Weight |
| --- | --- | --- | --- | --- | --- | --- |
| 7 | Location + month + line + location*month + location*line + month*line | 22 | -496.620 | 1037.2 | 0.00 | 0.504 |
| 11 | Location + month + line + location*month + location*line + month*line + sex | 23 | -496.217 | 1038.4 | 1.19 | 0.277 |
| 2 | Location + month + location*month | 12 | -508.137 | 1040.3 | 3.03 | 0.110 |
| 5 | Location + month + line + location*month | 14 | -506.842 | 1041.7 | 4.45 | 0.055 |
| 9 | Location + month + location*month + sex | 23 | -507.850 | 1041.7 | 4.46 | 0.054 |
| 4 | Location + month + line + month*line | 14 | -533.466 | 1094.9 | 57.69 | 0.000 |
| 6 | Location + month + line + location*line | 12 | -534.097 | 1096.2 | 58.95 | 0.000 |
| 3 | Location + month + line | 10 | -538.836 | 1097.7 | 60.43 | 0.000 |
| 10 | Location + month + line + sex | 11 | -538.836 | 1099.7 | 62.43 | 0.000 |
| 1 | location + month | 8 | -541.883 | 1099.8 | 62.53 | 0.000 |
| 8 | location + month + sex | 9 | -541.883 | 1101.8 | 64.53 | 0.000 |

To quantify the synchrony of oysters, we calculated the Shannon-Wiener Diversity Index (hereafter, diversity index) for each unique combination of month, location, and line (Shannon and Weaver 1949; Tuomisto 2012). The formula, where p_i_ is the proportion of the total sample found to have each of the five gametogenic categories, captures evenness of gametogenic stages at a given time and place (Eqn. 1). Used this way, the index has an inverse relationship with synchrony such that as the index increases, the level of synchrony decreases. The diversity index was compared across locations and lines using generalized linear models (Table 3), following the formulas created for models 1-7 of the 2018 ordinal logistic modeling (Table 2). Models were ranked in terms of AIC values, but because this diversity metric lacks replication within sites and lines, *post hoc* analyses were not possible. These analyses were performed using R and RStudio (R Core Team 2018).

**Table 3.**
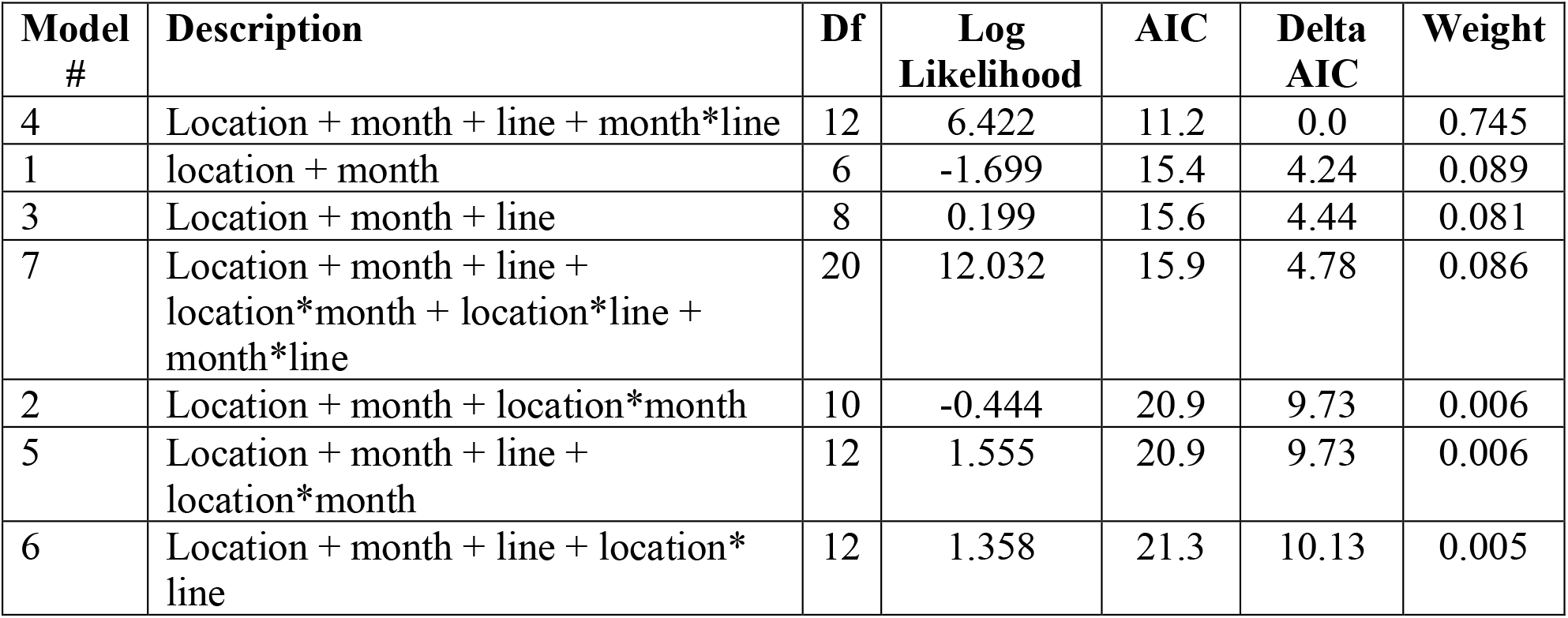
Results of 2018 model selection of generalized linear models of Shannon Diversity Index. Models are ranked in terms of AIC values

| Model # | Description | Df | Log Likelihood | AIC | Delta AIC | Weight |
| --- | --- | --- | --- | --- | --- | --- |
| 4 | Location + month + line + month*line | 12 | 6.422 | 11.2 | 0.0 | 0.745 |
| 1 | location + month | 6 | -1.699 | 15.4 | 4.24 | 0.089 |
| 3 | Location + month + line | 8 | 0.199 | 15.6 | 4.44 | 0.081 |
| 7 | Location + month + line + location*month + location*line + month*line | 20 | 12.032 | 15.9 | 4.78 | 0.086 |
| 2 | Location + month + location*month | 10 | -0.444 | 20.9 | 9.73 | 0.006 |
| 5 | Location + month + line + location*month | 12 | 1.555 | 20.9 | 9.73 | 0.006 |
| 6 | Location + month + line + location*line | 12 | 1.358 | 21.3 | 10.13 | 0.005 |

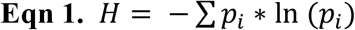

## RESULTS

### Environmental Conditions

Across locations, temperature varied from 0.9 - 30.1°C throughout the year (Fig. 2). There was a seasonal flux in temperature, with only slight variation between years and among locations. Treated as two regions, the northern sites (IRV, HH, SB) and southern sites (PGB, KCC) saw little difference in average temperature during any sampling month (Fig. 2). Salinity variation (Fig. 2) was non-overlapping for northern (0.9-15.7 psu) and southern sites (18.4-30.1 psu), supporting our use of these locations as proxy variables for low and high salinity, respectively. For each site, there was little variation in salinity between years, except at KCC and SB. In 2018, there was a gradual decrease in salinity at KCC (29.9 to 19.0 psu) between April and October, but in 2019 the trend was reversed, with salinity increasing from 23.8 to 30.1 psu in those months. The SB site had much lower spring salinity in 2019 than in 2018, but unfortunately insufficient oyster numbers remained at SB to allow an analysis for 2019.

**Fig. 2.**
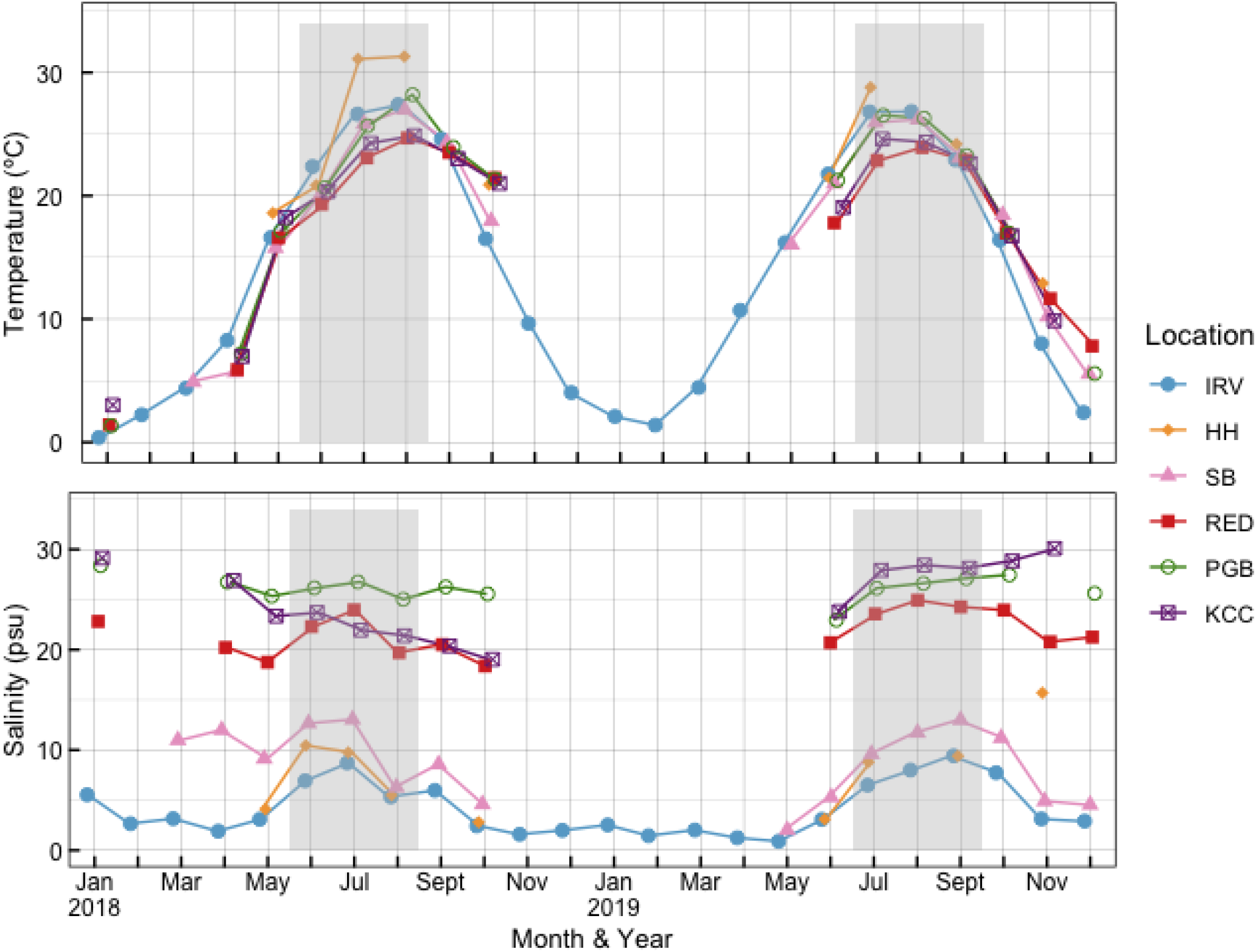
Temperature and salinity patterns in the Hudson River Estuary from January 2018 - December 2019. Shaded areas indicate sampling periods. Locations are listed in order by latitude, from north to south. Locality abbreviations explained in Fig. 1 legend and in the text

### Gametogenic Development Stage

Based on model selection criteria using AIC values, the top ranked model for 2018 was model 7: Location + month + line + location*month + location*line + month*line (Table 2). Model 11 was also supported, differing from model 7 only in the inclusion of sex as a parameter. Sex was determined to be an uninformative parameter based on the criteria in (Arnold 2010), so the *post hoc* analysis was thus performed only for model 7. All sampling occasions in 2018 and 2019 demonstrated either a roughly even or male-biased sex ratio (Table 4, Table S1). In August of 2018 and in all 2019 samples, the Aquaculture line at PGB became 100% male. For the linear models of the diversity index, the top model was model 4: location + month + line + month*line, indicating that location effects on synchrony were not interacting with line or month (Table 3).

**Table 4.** Sex ratios of oysters sampled for histology (# females: # males). Locations are listed by latitude, from north to south. Locality abbreviations explained in Fig. 1 legend and in the text. Native oysters in 2019 were dredged and outplanted as adults

| Location | Line | June 2018 | July 2018 | August 2018 | July/August 2019 | September 2019 |
| --- | --- | --- | --- | --- | --- | --- |
| IRV | Native | - | - | - | 11:9 | 11:9 |
| HH | Aquaculture | 3:10 | 7:13 | 7:13 | - | - |
|  | Hatchery | 3:10 | 5:7 | 4:8 | - | - |
|  | Native | 3:10 | 10:10 | 6:13 | 9:11 | 5:15 |
| SB | Aquaculture | 4:9 | 5:15 | 5:15 | - | - |
|  | Hatchery | 3:10 | 8:12 | 6:14 | - | - |
|  | Native | 2:11 | 7:13 | 7:13 | - | - |
| RED | Aquaculture | 2:8 | 1:19 | 2:18 | - | - |
|  | Hatchery | - | - | - | - | - |
|  | Native | - | - | 1:5 | - | - |
| PGB | Aquaculture | 2:11 | 5:15 | 0:20 | 0:20 | 0:20 |
|  | Hatchery | - | 6:12 | 4:16 | - | - |
|  | Native | 5:8 | 4:16 | 5:13 | 7:13 | 4:16 |
| **KCC** | Aquaculture | - | - | - | 4:16 | 7:13 |
|  | Native | - | - | - | 9:11 | 4:16 |

In June 2018, only Native and Aquaculture could be compared between low and high salinity sites and in both cases gametogenic development had advanced more at low salinity. A greater proportion of individuals were mature or spawning at the low salinity site SB (75-80%) compared to the high salinity site PGB (0-20%; Native p < 0.0001, n = 26; Aquaculture p = 0.0001 n = 26; Hatchery could not be tested; Tables 4, 5). For both lines, early and late active stages were found in no more than 20% of individuals at low salinity site SB, yet at high salinity these stages were still 100% and 70% of Native and Aquaculture samples, respectively (Fig. 3a, g). The low salinity site HH showed the same trend in both oyster lines, with a near absence of the early active stage, but was less extreme (post hoc contrast significant only for the Native line, Table 5). For the Native line at high salinity, the complete lack of mature or later stage individuals contrasted with the diverse gametogenic stages observed in Aquaculture at high salinity, while patterns were similar among all three lines at low salinity (Fig. 3). One exception is that Native and Hatchery lines achieved some June spawning at low salinity, but Aquaculture only had spawners at higher salinity sites in June. The diversity index was highest, and thus synchrony was lowest, for the Aquaculture line in the South (Table 6).

**Table 5.** Results of pairwise *post hoc* test (p-values) for the 2018 model comparing locations within each month, for each line. Locality abbreviations explained in Fig. 1 legend and in the text

|  |  |  |  |
| --- | --- | --- | --- |
| **a.) June** | Native | Hatchery | Aquaculture |
| HH-SB | 0.3761 | 0.0076 | 0.0347 |
| HH-PGB | 0.0002 | N/A | 0.1102 |
| SB-PGB | <.0001 | N/A | 0.0001 |
| **b.) July** | Native | Hatchery | Aquaculture |
| HH-SB | 0.4015 | 0.7865 | 0.9991 |
| HH-PGB | 0.4366 | 0.0581 | 0.0010 |
| SB-PGB | 0.0420 | 0.1531 | 0.0018 |
| **c.) August** | Native | Hatchery | Aquaculture |
| HH-SB | 0.9410 | 0.3073 | 0.6345 |
| HH-PGB | 0.0010 | <.0001 | <.0001 |
| SB-PGB | 0.0002 | 0.0026 | <.0001 |

**Table 6.** Shannon-Wiener Diversity Index values for each unique combination of month, location, and line for 2018. Locality abbreviations explained in Fig. 1 legend and in the text

|  | Location | Native | Hatchery | Aquaculture |
| --- | --- | --- | --- | --- |
| **June** | HH | 0.911 | 1.073 | 0.690 |
|  | SB | 0.937 | 0.790 | 0.687 |
|  | RED | - | - | 1.280 |
|  | PBG | 0.690 | - | 1.199 |
| **July** | HH | 0.562 | 0.888 | 0.999 |
|  | SB | 0.999 | 0.423 | 1.106 |
|  | RED | 0.673 | - | 1.476 |
|  | PBG | 1.141 | 0.557 | 1.234 |
| **August** | HH | 0.874 | 0.824 | 1.345 |
|  | SB | 0.708 | 0.824 | 1.345 |
|  | RED | - | - | 0.441 |
|  | PBG | 0.349 | 1.07 | 0.824 |

**Fig 3.**
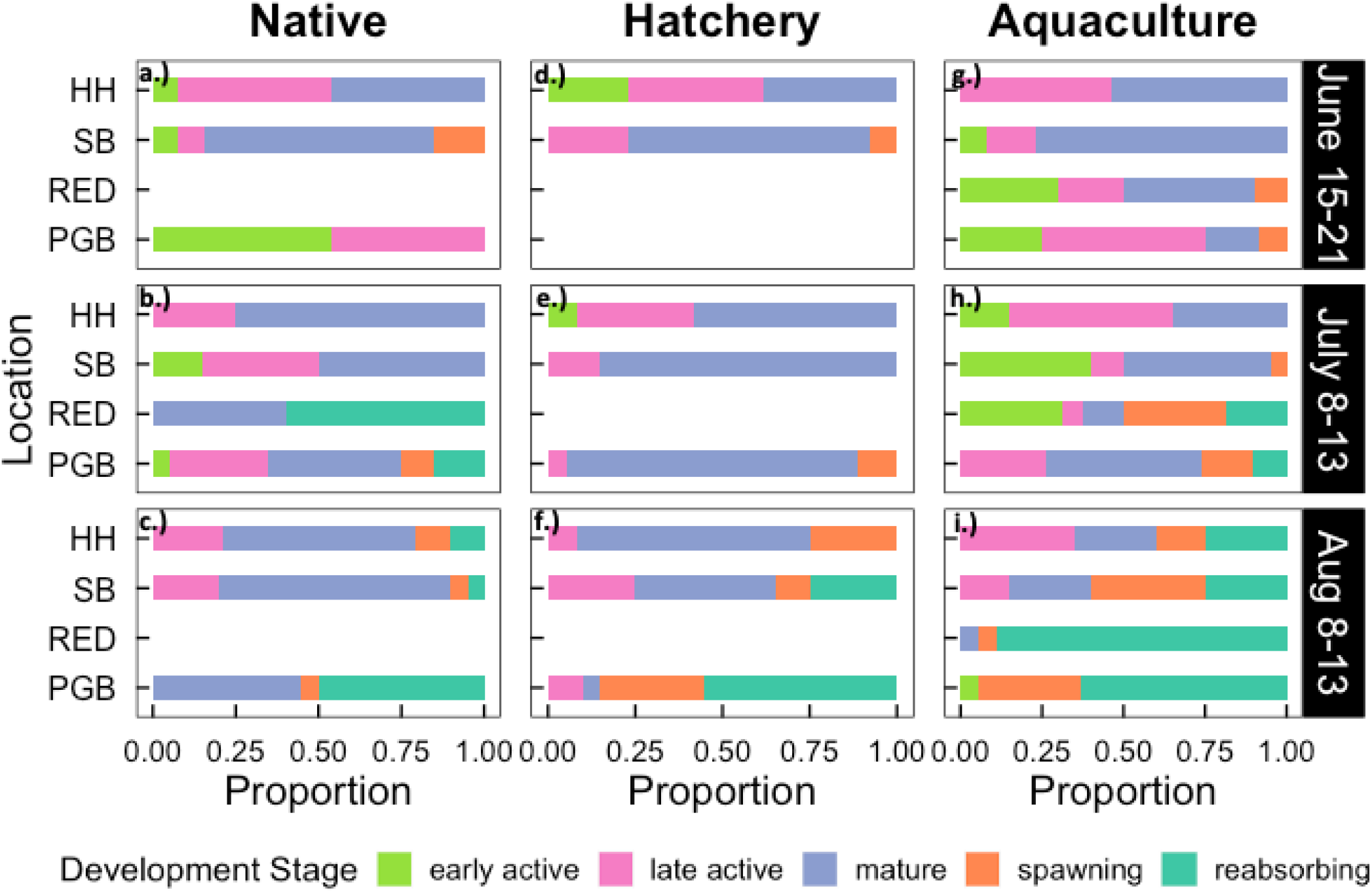
Gametogenic development stage proportions from 2018 oyster samples by month and location. a-c.) Native line. d-f.) Hatchery line. g-i.) Aquaculture line. Sample sizes were n=13 in June and n=20 in July and August with some exceptions (see Table 2). Locations are listed in order by latitude, from north to south. Locality abbreviations explained in Fig. 1 legend and in the text

In July, gametogenesis had not advanced much in any line at low salinity sites whereas at the moderate and higher salinity sites the proportion of spawning and reabsorbing stages had increased, where comparisons could be made to June. Thus, the June pattern of relatively advanced phenology at low salinity had reversed by July. Phenology again differed between low salinity SB+HH (average 0.9% spawning or reabsorbing individuals across the three lines and two sites) and high salinity PGB (21%), with 4 out of 6 post hoc contrasts p<0.05 (table 5). An even greater proportion of spawning and/or reabsorbing stages was observed at the moderate salinity RED site than at high salinity PGB. In contrast, the two low salinity sites, SB and HH, both were dominated by late active and mature stages with almost no spawners (Table 5, Fig. 3b, e, h). Comparing the ancestry lines, the most striking difference was a slightly slower phenology in Aquaculture as indicated by 18.6% early active oysters averaged across all sites compared with 4.5% in Hatchery and Native combined (Fig. 3b, e, h). In fact, Aquaculture phenology went slightly backwards from June to July (more early active stage at all sites except PGB), even while spawning increased (Fig. 3h). All ancestry lines contained all 5 stages when tallied over all locations in July, but the among-location average diversity index was 1.2 for the Aquaculture line, indicating lower synchrony than Native and Hatchery with 0.84 and 0.62 diversity indices, respectively (Table 6). The July Hatchery line within-locale synchrony was higher (diversity index lower) than in any other month and line (Table 6).

In August, gametogenic progression to a later mix of stages was observed in all lines, at all sites. All three lines continued to show a higher proportion of later stages at higher salinity sites, dominated by spawning and reabsorption, relative to low salinity sites where late-active and mature stages were abundant (all p<0.0026, Table 5, n=51, 60, 58, for HH, SB, PGB, respectively all n=20; Table 4). Relative to July, low salinity sites in August progressed less than higher salinity sites except for the Aquaculture line. The only other line distinction of note was the Native oysters having 44% pre-spawning stages at high salinity PGB, whereas Hatchery and Aquaculture lines were >85% spawning and reabsorbing stages (Fig. 3 c, f, i). The diversity index was again overall highest for the Aquaculture line.

The 2019 data only included two months, and data collection was shifted 2 weeks later relative to 2018. Two sites were included for both the high salinity and low salinity regions, but complete data were only obtained for the Native line (adult transplants that had acclimated for a year in this case, as opposed to 2 years growth from spat on site for 2018 data). No Hatchery line oysters were available in 2019. At all sites oysters demonstrated gametogenic progression between the two sampling dates (Fig. 4). The *post hoc* contrasts between locations found little difference in stage proportions for either month (Table 7, Fig. 4a, b). This spatial gametogenic synchrony was coupled with temporal synchrony within sites based on low average diversity index values of 0.38 and 0.44 for July/August and early September, respectively (Table 8). At the high salinity sites where the Aquaculture and Native lines could be compared, the former had substantially more advanced gametogenic stages at the end of July (90+% spawning and reabsorption stage for Aquaculture, but only 15-45% for Native), but only modest differences with the same trend at the beginning of September.

**Table 7.**
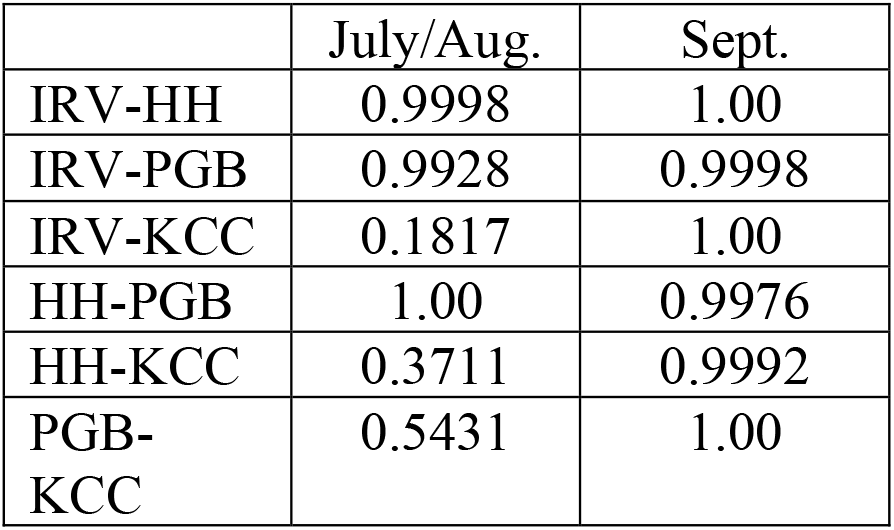
Results of pairwise *post hoc* test (p-values) for the 2019 model comparing locations within each month for the Native line. Locality abbreviations explained in Fig. 1 legend and in the text

**Table 8.** Shannon-Wiener Diversity Index values for each unique combination of month, location, and line for 2019. Locality abbreviations explained in Fig. 1 legend and in the text. *Native oysters dredged and outplanted as adults

|  | Location | Native* | Aquaculture |
| --- | --- | --- | --- |
| **July/Aug** | IRV | 0.199 | - |
|  | HH | 0.381 | - |
|  | PGB | 0.263 | 0.525 |
|  | KCC | 0.680 | 0.276 |
| **Sept** | IRV | 0.491 | - |
|  | HH | 0.514 | - |
|  | PGB | 0.376 | 0.263 |
|  | KCC | 0.387 | 0 |

**Fig 4.**
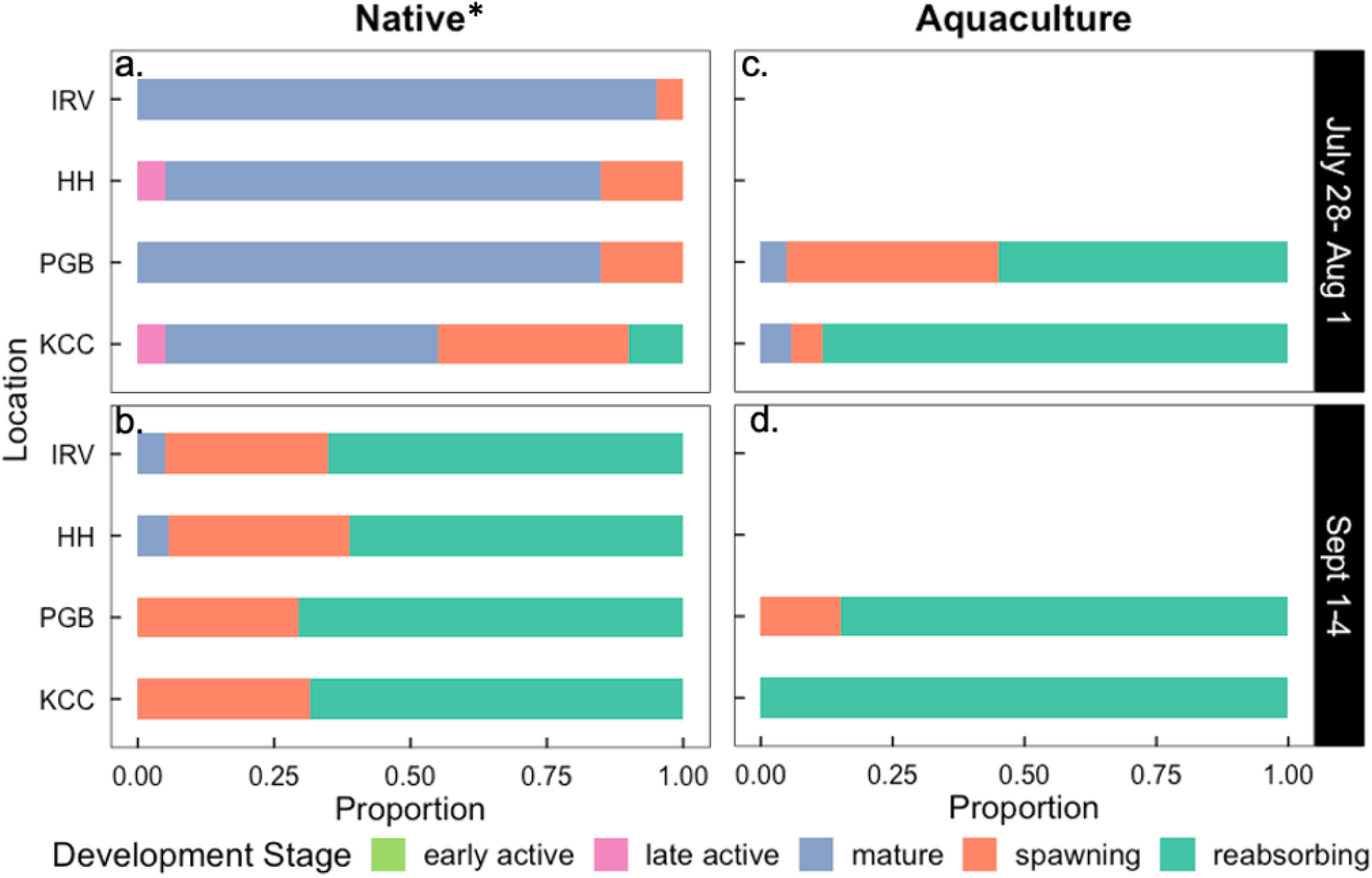
Gametogenic development stage proportions for 2019 oyster samples by month and line, including Native line (a-b.) and Aquaculture line (c-d.). N=20 in all cases. Locations are listed in order by latitude, from north to south. Locality abbreviations explained in Fig. 1 legend and in the text. *Native oysters dredged and outplanted as adults

### Dermo Disease

Only 3 of 240 individuals had *P. marinus* spores detected (1.3%). All detected infections were in the Native line at PGB during the September sampling period. Two samples had Mackin intensity levels of 0.5, and another had a level of 1.

## DISCUSSION

In one of the first studies to measure salinity effects on gametogenic phenology in a bivalve, and compare oyster lines with different ancestries, we found that in July and August of 2018 all three oyster lines had delayed gametogenic development at low salinity sites relative to high salinity sites, confirming our first hypothesis that low salinity would cause a delay.

Surprisingly, the reverse was true in June 2018 for the two lines that could be compared in different salinities up and down river. June spawners only occurred at low salinity for Native oysters and contributed to a weighted average gonad index of 2.38-2.92 at low salinity sites, compared to only 1.46 at high salinity. The Aquaculture line showed this trend less dramatically, with weighted averages of 2.62 vs. 2.08 at low and high salinity, respectively, partly because the few June spawners were observed only at high and moderate salinity sites. The June pattern of more advanced oyster gonad index proportions at low salinity relative to high salinity was reversed in July due to gonad index stasis at low salinity versus progression toward spawning and reabsorption at high salinity.

Our second hypothesis, that the Native line would have the most distinct phenology characterized by earlier (less delayed) gametogenesis at low salinity, was rejected. All three lines had similar gonad index distributions at low salinity in June. Instead, the most striking distinction observed in the Native line was the extent of gonad index delay at high salinity in June, as indicated by zero oysters with a ‘maturing’ or later stage (weighted average gonad index 1.46), whereas at the same site the Aquaculture line had 25% maturing or spawning oysters (2.08). By July the Native oysters had compensated and had gonad index proportions similar to the other lines. The other line distinction of note in 2018 was the consistently higher diversity of gonad index stages in the Aquaculture line throughout the summer, indicating 30-48% less synchrony at any particular time or place.

Low variation in temperature among sites and minimal Dermo infections detected overall, highlight regional salinity differences as likely key to phenology variation among sites in 2018, but causal inferences are tentative because oyster cage outplant sites differed in other unmeasured ways. The 2019 sampling provided fewer spatial and line comparisons. Despite similar environmental conditions in the two years, 2019 patterns indicated greater synchrony and only slight salinity effects relative to 2018. We discuss these findings with reference to environmental variation and future restoration goals.

### Phenology variation

#### Month

The model that best explains 2018 gametogenic stage proportions includes month, location, line, and their interactions. Given the summer temperature variation (Fig. 2), month is the most obvious factor of expected importance since the species is considered to have locally synchronous spawning triggered by rising temperatures and conspecific spawning (Thompson et al. 1996; Bernard et al. 2011; Aranda et al. 2014). While some spawning occurred in each month (average proportion of spawners and reabsorbing individuals equal to 3.5%, 7.5%, and 46.8% in June, July, and August 2018, and 58.63% and 97.5% in July/August and September 2019), the spawning period in the HRE peaked in August and continued into September for these outplanted, caged oysters. An August peak for native TZHB oysters is supported by spat monitoring in the HRE, where the most abundant recruitment of spat (after a 2-3 week larval stage) has been observed in September (McFarland and Hare 2018). Our results suggest an extension of the spawning season for the HRE relative to nearby Long Island Sound where spawning peaks in July with spent oysters by August (Ford et al. 1990; Barber et al. 1991). Cold Adirondack mountain Spring melt waters in the Hudson River, or differences in phytoplankton bloom timing and concentration, could be responsible for this geographic difference (Hofmann et al. 1992; Bernard et al. 2011).

#### Salinity

Our hypothesis of delayed gametogenic progression at lower salinities was only observed after the month of June, suggesting that the initiation of gametogenesis is less constrained by salinity variation, but that low summer salinity at northern sites relative to southern sites could be causing a delay at later gonad maturation stages. By the August 8-13 sampling period in 2018, northern low salinity sites (HH, SB) had fallen to near 5 psu and a majority of oysters were still in pre-spawning stages, a trend we expected based on literature suggesting reproductive depression caused by low salinity (Butler 1949; Shafee and Daoudi 1991; Shumway 1996; Honig et al. 2014; Volety et al. 2017). In contrast, the majority of oysters at the high salinity PGB site (approximately 25 psu) were spawned or reabsorbing for all three lines in August. These June and later Summer findings in Hudson River oysters are similar to those reported by McFarland et al. (2022) for eastern oysters in the Caloosahatchee River Estuary in Florida. They found over 15 years of monthly sampling, across 5 sites ranging from 12 to 28 average salinity, that average gonad index at sites had a negative relationship with salinity. The lowest salinity site had the earliest gametogenic development; it had a 15-year average March gonad index of 2 while all other sites had averages below 1.5 (Fig. 7 in McFarland et al. 2022). We did not measure Spring gametogenesis in Hudson River oysters, but early initiation in the northern TZHB population seems unlikely given that it often experiences extended periods below salinity 5 during Spring (Levinton et al. 2011). Instead, we speculate that the advanced June gametogenesis observed here results from a rapid compensation for stressful Spring conditions. Whatever the mechanism, the early gametogenic progression that produced some spawning in northern low salinity sites, coupled with a small proportion of spawners in August relative to high salinity sites, indicates that oysters at low salinity sites in the Hudson River also had relatively extended reproductive seasons similar to what was reported in the Caloosahatchee River (McFarland et al. 2022).

Contrary to salinity effects observed in 2018, oysters analyzed in 2019 showed no significant differences in gonad index among sites. We don’t know what interannual differences caused this change in phenology pattern along the salinity gradient, but the summer variation in salinity at northern sites was less in 2018, lacking a 5 psu drop in August or September (Fig. 2). Phenology comparisons between years are tenuous with respect to salinity contrasts because of the varied sampling dates, uneven oyster line availability, and altered Native outplant method. Factors such as food availability (Luna et al. 2000; Bernard et al. 2011) or local freshwater inputs (i.e. river discharge) (Baba et al. 1999; Wilson et al. 2005) could drive interannual variation, but such factors were outside the scope of this study. We recommend further study to determine the key interannual factors that can change reproductive phenology patterns so dramatically.

In terms of outplant method, it is notable that the 2019 Native dredged oysters were able to develop gametes at a variety of salinities only one year after transplant. Furthermore, the fact that both outplant strategies (outplanted as spat, outplanted dredged adults) resulted in oysters that were able to undergo a full reproductive cycle is encouraging for future restoration efforts considering the potential advantages associated with restoration using locally-adapted species (Hofmann et al. 1992; Camara and Vadopalas 2009; Flanagan et al. 2018; Hornick and Plough 2019; Hornick and Plough 2022). This phenotypic plasticity is characteristic of oysters, but presumably has limits that need to be accounted for in restoration planning (Li et al. 2017; Li et al. 2018; Li et al. 2021).

#### Ancestry

The prediction that the Native line is adapted to low salinity and therefore would uniquely demonstrate a lack of reproductive depression at low salinity was only consistent with patterns observed in June 2018. Unfortunately, only comparisons between Native and Aquaculture strains were possible in June. The spatial contrast in gonad index between low and high salinity sites was much stronger in the Native line. The average proportion of early active and mature individuals for Natives in the north was 0.08 and 0.58 respectively, compared to 0.54 and 0.0 at high salinity PGB. The overall gonad index average for Natives was higher at low salinity vs high salinity sites, 2.65 vs 1.46. This trend also was detectable in Aquaculture individuals but was less extreme (gonad index 2.62 in the north vs 2.08 at high salinity PGB).

This suggests that TZHB Native oysters may have local adaptations that allow them to rapidly compensate for gametogenic delays experienced during very low Spring salinity levels, perhaps in response to overall metabolic depression (Gurr et al. 2020).

In August there was no single oyster line that stood out in terms of phenology or salinity effects, but there were differences in the gonad index progression with Native slowest (3.37 average gonad index), Hatchery intermediate (3.54) and Aquaculture most advanced (3.82).

These differences and trend are similar in the northern low salinity sites vs high salinity PGB, just shifted lower and higher than the overall average, respectively (3.03, 3.17, 3.50 in north; 4.06, 4.30, 4.45 at PGB). This trend for 2018 indicates that under some conditions the Native line has a more extended reproductive season than the other oyster lines. A lengthening of the gametogenic period can be interpreted as a depression of gametogenesis that has been observed by previous studies. (Butler 1949; Shafee and Daoudi 1991; Shumway 1996; Honig et al. 2014; Volety et al. 2017).

The 2019 samples allowed line comparisons only at high and low salinity sites. At these sites the gonad index differences between lines were not large, but the 2018 trend of a more extended reproductive season for the Native line was repeated in 2019. At the end of July gonad index averages for low and high salinity were 3.05 and 3.85 for Native, and 4.66 at high salinity sites for Aquaculture. In early September the difference was similar, 4.56 and 4.70 for Native, and 4.93 at high salinity sites for Aquaculture. In September, the transplanted Natives were majority post-spawning at low salinity sites, suggesting that the Natives had completed a cycle of gametogenesis and spawning.

One of the biggest differences among lines was in synchrony as measured by the diversity index. When averaged across all sites and over the entire 2018 season, the stage diversity in Native, Hatchery, and Aquaculture was 0.78, 0.81, and 1.05, respectively. Thus, Aquaculture was the outlier with less synchrony. We are not aware of other comparable data and can only speculate that long term culture and domestication led to relaxed selection (i.e., more random change under genetic drift) on sensory systems that contribute to synchrony under natural conditions. However, the same strain difference in synchrony was not observed in 2019, so this difference among strains probably relates to distinct expressions of genotype by environment plasticity (Eierman and Hare 2015).

#### Conclusions and context for future oyster restoration

This study demonstrated differences in reproductive phenology along the HRE salinity gradient that were associated with locality (low vs. high salinity regions) and oyster ancestry line. These novel results in a temperate estuary suggest that reproductive phenology within an eastern oyster metapopulation is not simply a function of temperature, but also varies along the salinity gradient. The 2018 patterns were consistent with salinity effects on phenology documented elsewhere, but our interannual results caution against generalizing from the 2018 patterns because genotype by environment interactions likely have strong effects. Additional replication is necessary to further understand genotype by environment effects. Future studies would be valuable that can experimentally link aspects of phenology with specific environmental drivers. In particular, future studies should sample in Spring as well as Summer because salinity effects seem to vary at different stages, and experimental designs to test for reproductive compensation would help evaluate the consequences of stressors.

Similar to previous reports in the HRE (Levinton et al. 2011; McFarland and Hare 2018), the near absence of Dermo infections in this study makes it reasonable to infer salinity effects separate from disease interactions. Further, the lack of Dermo presence across all sites supports the conclusion that disease is not the reason for the Native TZHB population’s current isolation at low salinities, but further study would be necessary to confirm this.

The moderate salinity Aquaculture line studied here succeeded in spawning across diverse environments, attesting to its plasticity with respect to salinity. However, it also showed lower gametogenic within-site synchrony compared to the other lines, perhaps indicating domestication effects lowering a fitness-related trait. Selective breeding for commercially valuable traits can cause reduced genetic diversity and potentially reduced lifetime fitness in a natural environment, making aquaculture lines often the least desirable option for restoring oyster populations (Baggett et al. 2014). Enhancing or restoring a population with stocks that do not possess adequate genetic variation, or that have maladaptive variation, could be detrimental to overall population fitness and diversity (Camara and Vadopalas 2009; Morvezen et al. 2016; Hornick and Plough 2019; Hughes et al. 2019; Hornick and Plough 2022).

With the growing threat of climate change, species inhabiting estuaries will likely be exposed to more frequent and extreme storms that can rapidly shift salinity to the edge of their tolerances. Investigating the effect of salinity on reprodutive phenology is only one aspect of research that is essential to understand and possibly mitigate the potential effects of climate change on estuarine oyster populations. The Native TZHB population is at the edge of habitable salinity variation with little quality habitat to expand to upstream (Starke et al. 2011), and it is not known to occur in any habitat further south except as sparse spat (new recruits; Medley 2010) or adults on pilings in Hudson River Park, Manhattan (Fitzgerald et al. 2020). The ability of TZHB oysters to survive and reproduce at moderate salinities, as evidenced in this study, indicates that its continued isolation north of New York City is likely a function of hydrodynamics and/or stressful downstream water quality for the mobile larval stage. If poor water quality in the lower HRE amplifies larval mortality above typical levels, then further water quality improvements will be needed before restoration with any oyster line has a chance of establishing a self-sustaining metapopulation (i.e., promoting the full life cycle) in the lower estuary. Assuming adequate water quality, transplantation of adults or spat to seed hard substrates in moderate salinity habitats could help protect the population from climate change effects by reestablishing metapopulation dynamics (Lipcius et al. 2008). Here, results demonstrate that the TZHB population provides a valuable resource for transplant restoration strategies that could leverage and expand local genetic variation.

## ACKNOWLEDGEMENTS

Thanks to Harmony Borchardt-Wier, Iris Burt, Sarah Gisler, Hannah Hartung, and Tiffany Medley for their support in this research effort through field work assistance and/or methods and data analysis consults. We appreciate the careful oyster husbandry contributed by Amandine Hall at Martha’s Vineyard Shellfish Group and by Gregg Rivara at the Suffolk County Marine Environmental Learning Center hatchery. Pete Malinowski generously provided some of the oysters analyzed in this study. Thanks to Captain Jim Nickels, Monmouth University, for a successful oyster dredging trip in 2018, ultimately allowing for this experiment to continue into 2019. Advice provided by Cornell Statistical Consulting Unit on data analysis and display was greatly appreciated. This study would not have been possible without housing and lab space provided by the U.S. Forest Service Fort Totten Urban Field Station. Funding was provided by a Tibor T. Polgar Fellowship from the Hudson River Foundation, a grant from the NYS Water Resources Institute, and a grant from the Cornell University College of Agriculture and Life Sciences Charitable Trust. This work was only possible through the generosity of location hosts that kept experimental oyster cages secure: Irvington Boat Club, Hastings on Hudson Neighborhood Association, Groundwork Hudson Valley Science Barge, Inwood Canoe Club, Red Hook Barge Museum, Sebago Canoe Club, and Kingsborough Community College.

## ELECTRONIC SUPPLEMENTARY INFORMATION

**Table S1.** Lengths of oysters sampled for histology by sex (female, male, average length (mm) *±* S.E). Locations are listed in order by latitude, from north to south. Locality abbreviations explained in Fig. 1 legend and in the text

| Location | Line | June 2018 | July 2018 | August 2018 | July/August 2019 | September 2019 |
| --- | --- | --- | --- | --- | --- | --- |
| **IRV** | Native | - | - | - | 69.0 ± 4.5:<br>66.4 ± 4.9 | 71.2 ± 4.4,<br>64.6 ± 4.8 |
| **HH** | Aquaculture | 36.7 ± 1.8,<br>32.9 ± 2.2 | 45.2 ± 3.8,<br>41.0 ± 2.3 | 51.4 ± 4.2,<br>49.0 ± 2.5 | - | - |
| | Hatchery | 53.3 ± 3.8,<br>46.5 ± 1.8 | 35.6 ± 2.7,<br>34.6 ± 2.4 | N/A | - | - |
| | Native | 34.7 ± 4.5,<br>36.6 ± 1.3 | 42.8 ± 3.8,<br>38.7 ± 3.1 | N/A | 63.9 ± 3.4,<br>62.6 ± 5.1 | 71.2 ± 4.4,<br>64.6 ± 4.8 |
| **SB** | Aquaculture | 63.0 ± 7.4,<br>55.9 ± 4.8 | 45.8 ± 3.0,<br>45.5 ± 1.6 | 53.7 ± 5.6,<br>51.6 ± 4.1 | - | - |
| | Hatchery | 52.7 ± 3.4,<br>49.1 ± 2.7 | 56.0 ± 6.4,<br>59.0 ± 5.6 | 74.0 ± 3.6,<br>61.5 ± 2.1 | - | - |
| | Native | 42.0 ± 0.0,<br>38.2 ± 2.3 | 49.7 ± 3.2,<br>42.0 ± 2.1 | 47.1 ± 3.1,<br>52.0 ± 2.4 | - | - |
| **RED** | Aquaculture | 70.5 ± 17.5:<br>60.5 ± 6.2 | 64.0 ± 0,<br>45.6 ± 4.2 | 72.0 ± 8.0,<br>57.8 ± 3.6 | - | - |
|  | Hatchery | - | - | - | - | - |
| | Native | - | - | 45.0 ± 0,<br>48.2 ± 2.6 | - | - |
| **PGB** | Aquaculture | 94.5 ± 1.5,<br>68.5 ± 4.3 | 55.2 ± 3.0,<br>59.4 ± 2.9 | 0,<br>52.7 ± 2.7 | 0,<br>95.0 ± 2.4 | 0,<br>91.3 ± 2.7 |
| | Hatchery | - | 65.3 ± 3.1,<br>57.8 ± 3.0 | 48.2 ± 5.0,<br>58.3 ± 3.2 | - | - |
| | Native | 49.2 ± 3.0,<br>44.6 ± 1.7 | 45.5 ± 3.5,<br>43.7 ± 1.7 | 45.8 ± 3.1,<br>52.8 ± 3.0 | 67.1 ± 4.7,<br>68.6 ± 3.2 | 81.5 ± 8.2,<br>68.3 ± 2.6 |
| **KCC** | Aquaculture | - | - | - | 91.5 ± 8.0,<br>85.7 ± 3.4 | 91.0 ± 5.8,<br>82.5 ± 3.7 |
| | Native | - | - | - | 73.1 ± 5.6,<br>70.6 ± 4.6 | 86.5 ± 4.4,<br>76.9 ± 2.7 |

